# Isolation and characterization of SARS-CoV-2 VOC, 20H/501Y.V2, from UAE travelers

**DOI:** 10.1101/2021.05.14.443968

**Authors:** Pragya D. Yadav, Prasad Sarkale, Alpana Razdan, Nivedita Gupta, Dimpal A. Nyayanit, Rima R. Sahay, Varsha Potdar, Deepak Y. Patil, Shreekant Baradkar, Abhinendra Kumar, Neeraj Aggarwal, Anita M. Shete, Harmanmeet Kaur

**Author notes:** **Corresponding author**, Dr. Pragya D. Yadav, Scientist ‘E’ and Group Leader, Maximum Containment Facility, Indian Council of Medical Research-National Institute of Virology, Sus Road, Pashan, Pune, Maharashtra, India Pin-411021.

## Abstract

Multiple SARS-CoV-2 variants have been emerged and created serious public health in the affected countries. The variant of Concern associated with high transmissibility, disease severity and escape mutations is threat to vaccination program across the globe. Travel has been important factor in spread of SARS-CoV-2 variants worldwide. India has also witnessed the dreadful effect of these SARS-CoV-2 variants. Here, we report the Isolation and characterization of SARS-CoV-2 VOC, 20H/501Y.V2 (B.1.351), from UAE travelers to India. The virus isolate would be useful to determine the efficacy of the currently available vaccines in India.

## Introduction

Since its emergence in 2019, SARS-CoV-2 has effectively evolved accumulating the deleterious mutations in its genome. Within the last five months, multiple SARS-CoV-2 variants have been detected across the globe which are more transmissible, possibly escapes the natural and vaccine-induced immunity, and could lead to increased SARS-CoV-2 infection [1, 2]. Of these, SARS-CoV-2 20H/501Y.V2 (B.1.351) variant is the most dreadful variant reported to emerge from South Africa in December 2020; however, its earliest detection was traced back to October 2020. It has been now the most prevalent lineage in South Africa and has also been reported from 68 countries. B.1.351 variant has 21 mutations with 9 spike protein mutations in the genome. The key mutations beyond N501Y are E484K, K417N, orf1b deletion in the Receptor binding domain (RBD) and L18F, D80A, D215G, Δ242-244, R264I, A701V in the N terminal domain [3]. Pearson et al. had estimated that the B.1.351 variant could be highly transmissible than the earlier circulating strains of SARS-CoV-2. It has been observed that B.1.351 accounted for about 40% of new SARS-CoV-2 infections compared to only 20% for B.1.1.7 variant. An in vitro study on the monoclonal antibodies and convalescent plasma samples of COVID-19 cases demonstrated reduced activity against B.1.351 compared to B.1.1.7 [4]. The findings of the NVX-CoV2373 clinical trial have shown post hoc vaccine efficacy of 51% in South Africa where B.1351 was prevalent [5]. The ChAdOx1 nCoV-19 clinical trials results didn’t show protection against mild-moderate infection with B.1.351 [6]. Besides this, B.1.351 is also reported to be less susceptible to the currently available vaccines i.e., ChAdOx1 nCoV-19, mRNA-1273, BNT162b2, NVX-CoV2373 [6–10]. It is presumed that natural and vaccine-induced immunity would not protect against B.1.351.

With its worldwide occurrence, B.1.351 has been an addition to the gruesome situation of the SARS-CoV-2 pandemic. India has also reported the presence of various SARS-CoV-2 variants such as B.1, B.1.1.7, B.1.1.28.2, and B.1.617.1 [11–14]. A continuous effort of virus isolation of VOC B.1.351 was made from COVID-19 positive individuals who traveled from foreign countries to India. Here, we report the isolation and characterization of VOC B.1.351 from the foreign travelers who arrived in India.

## Methods

### Clinical specimens

With ongoing process of SARS-CoV-2 screening of travelers from foreign countries to India, the oropharyngeal and nasopharyngeal swab specimens were collected from 58 individuals with travel history from United Arab Emirates (UAE) (n=39), East and West Africa (n=10), Qatar (n=5), Ukraine (n=3) and Saudi Arabia (n=1) arrived at New Delhi International airport in India. All the subjects were asymptomatic and found to be SARS-CoV-2 positive by real time RT-PCR (Supplementary table 1) [15, 16].

### Virus isolation and titration

Vero CCL-81 cells were grown to confluent monolayer in 24-well plate maintained in Eagle’s Minimum essential medium (MEM) supplemented with 10 %/ fetal bovine serum (FBS) (HiMedia, Mumbai), penicillin (100 U/ml) and streptomycin (100 mg/ml). After decanting the growth medium, one hundred microliter volume of clinical specimens of 58 subjects were inoculated onto 24-well cell culture monolayer of Vero CCL-81. The cells were incubated for one hour at 37°C to allow virus adsorption, with rocking every 10 min for uniform inoculum distribution. After the incubation, the inoculum was removed and the cells were washed with 1× phosphate-buffered saline (PBS). The MEM supplemented with two percent FBS was added to each well. The culture was incubated further in incubator at 37°C with 5% CO_2_ and observed daily for cytopathic effects (CPEs) under an inverted microscope (Nikon, Eclipse Ti, Japan) [11]. The virus titration was carried out using the cell culture supernatants of the clinical specimens displaying CPE in cell culture. Median tissue culture infective dose (TCID_50_) value was calculated using the Reed and Muench method [17].

### Genomic characterization of SARS-CoV-2 isolates

Genomic characterization of the virus isolates was carried out using Next-Generation Sequencing (NGS) with the quantified RNA. Briefly, the ribosomal RNA depletion was performed using Nebnext rRNA depletion kit (Human/mouse/rat) followed by cDNA synthesis using the first strand and second synthesis kit. The RNA libraries were prepared using TruSeq Stranded Total RNA library preparation kit. The amplified RNA libraries were quantified and loaded on the Illumina sequencing platform after normalization [18]. The SARS-CoV-2 sequences were retrieved using the reference-based assembly method and Wuhan Hu-1 as the reference sequence (NC_045512).

## Results

### Virus isolation and titration

Cytopathic effect (CPE) was observed in four out of fifty-eight specimens with the syncytial formation on post-infection day (PID)-2. The progressive infectivity was observed with the fusion of the infected cells with neighboring cells leading to the generation of the large mass of cells on 3rd PID (Figure 1 A). The presence of the replication-competent virus was confirmed by Real-time RT-PCR that demonstrated a higher viral load of 4.6×10^9^ to 1.8×101^0^ and 8.4×10^11^ to 1.5×10^12^ in the cell culture medium for 2nd and 3rd post infection day than inoculated specimens [15, 16]. Culture that shown CPE were centrifuged at 4815 × g for 10 min at 4°C; the supernatants were processed immediately or stored at −80°C The four virus isolates titrated at passage 3 demonstrated the virus titer of 10^5.5^⍰10^5.66^TCID50/ml respectively.

**Figure 1.**
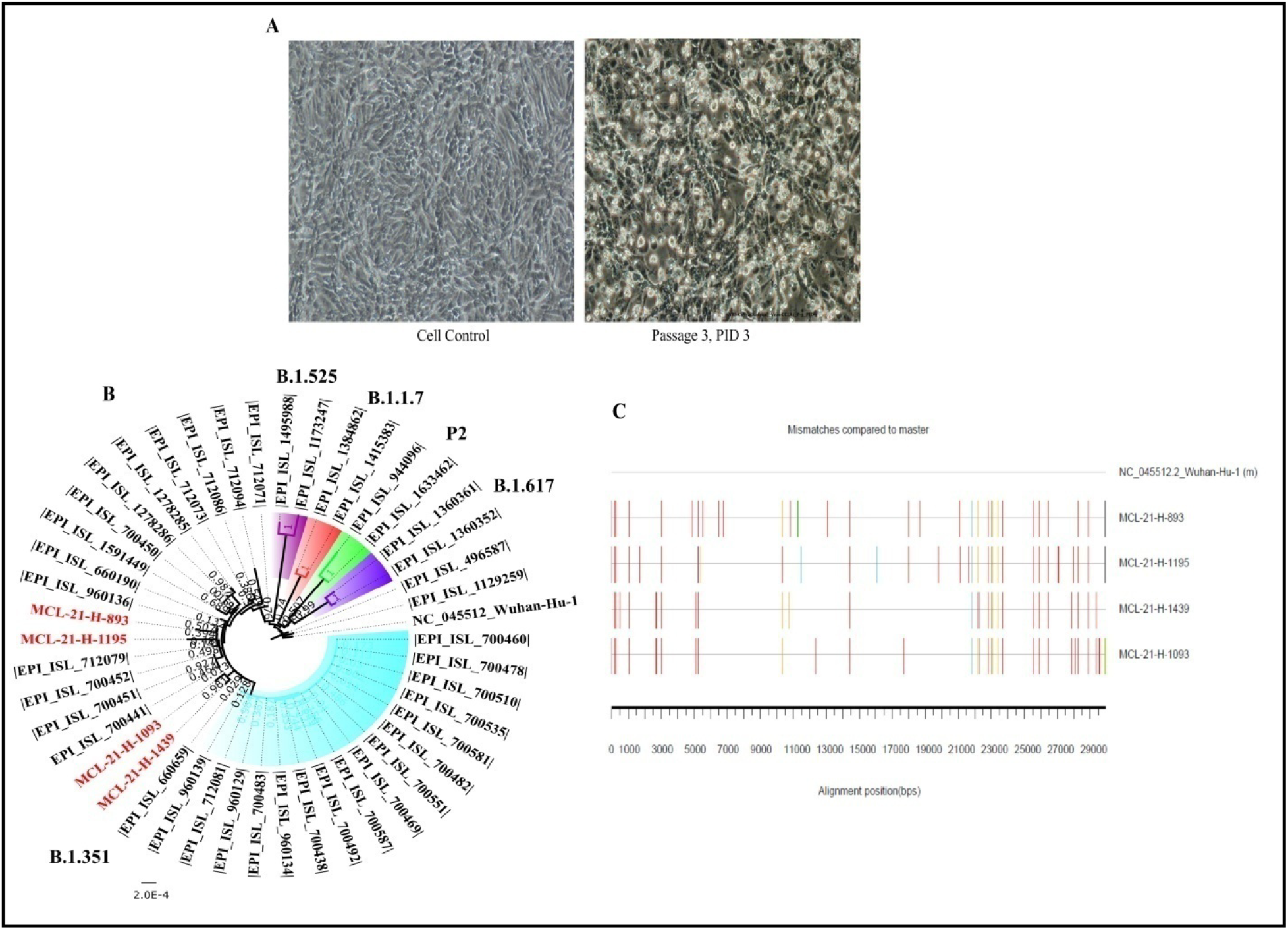
Isolation and bioinformatics analysis of B.1.351 SARS CoV-2 sequences: A) Cytopathic effect observed on the Vero CCL81 cell line for the 3^rd^ passage at 3^rd^ PID along with the control (left). B) Neighbor joining tree of the isolate sequences with reference sequences downloaded from GISAID with Tamura’s 3 Parameter model and a bootstrap replication of 1000 cycles. The sequences retrieved in study are colored red. C) The in B.1.351 SARS CoV-2 sequences retrieved in this study are aligned with reference isolate of Wuhan-HU-1 (Accession No.: NC_045512.2). The nucleotide changes are generated using the highlighter plot available at https://www.hiv.lanl.gov/content/sequence/HIGHLIGHT/highlighter_top.htm. The mismatches are marked in different colors adenine (A) green; cytosine (C) blue; guanine (G) orange, thymine (T) red and gap black color.

### Clinical history of the cases

All four cases were male, asymptomatic, and belonged to the age group of 27, 32, 42, and 68 years (Supplementary table 1). None had any co-morbid conditions. All these cases recovered completely with no new symptoms or complications. Currently, there is no evidence to suggest that this variant has caused any disease severity in these cases.

### Genomic characterization of SARS-CoV-2 isolates

The neighbor-joining tree demonstrated a separate cluster each consisting of two SARS-CoV-2 sequences; cluster1: MCL-21-H-1195 (EPI_ISL_2014135) and MCL-21-H-893 (EPI_ISL_2014131) and cluster 2: MCL-21-H-1093 (EPI_ISL_2014132) and MCL-21-H-1439 (EPI_ISL_2014133) (Figure 1B). EPI_ISL_2014132 and EPI_ISL_2014133 shared common mutations at genomic positions A2692T, C5100T, G27870T, and C29358T. Figure 1C depicts the nucleotide mismatches observed in the isolates. All the four SARS CoV-2 isolate sequences retrieved had common spike mutation (D80A, D215G, L242_L244del, K417N, E484K, N501Y, D614G and A701V) (Supplementary table 2). These isolates also have common mutations in the 5’ UTR (G174T, C241T), ORF1ab (C3037T) and ORF8 (28253). The L18F and R246I amino acid mutations described by Tegally et al [3] were absent in the spike protein in each isolate (Supplementary table 2).

The sequence analysis of South African B.1.351 lineage sequences demonstrated the presence of two different clusters having a deletion of nine nucleotides each in the ORF1ab (GP:11288-11296) and the spike (GP:22287-22295) while the second clusters with deletion of nine nucleotides in spike (GP:22287-22295) region. The B.1.351 lineage isolates retrieved in this study have nine nucleotides deletion in both ORF1ab and spike region. The percent nucleotide similarity demonstrated a 99.88-99.96% similarity of the isolates with the representative B.1.351 lineage sequences (Supplementary table 3).

## Discussion

The global increase in the number of SARS-CoV-2 infections necessitates the rigorous implementation of the vaccination program and intervention measures such as hand hygiene, masking, physical distancing and restrictions on public gatherings to curb the spread of SARS-CoV-2. Besides this, the emergence of these new variants has been a serious threat to the COVID-19 vaccination program. Hence, it is necessary to determine the neutralizing capacity of the newly emerging SARS-CoV-2 VUI and VOC variants against the available vaccines. The isolation of B.1351 would be useful to assess the efficacy of the vaccines rolled out under the national COVID-19 vaccination program of India and also developing new vaccine, diagnostic or antiviral testing if needed and situation demand in future.

## Ethical approval

The study was approved by the Institutional Biosafety Committee and Institutional Human Ethics Committee of ICMR-NIV, Pune, India under project ‘Propagation of new SRS-CoV-2 variant isolate and characterization in cell culture and animal model.

## Author contributions

PDY and NG contributed to study design, data analysis, writing and critical review. PS, SP, SB and AK performed the laboratory experiments, interpretation, and data analysis. AS, RRS, DAN, DYP, HK, NA, VP and AR contributed to data collection, interpretation, writing and critical review.

## Acknowledgement

The author gratefully acknowledges the contribution of Mr. Rajen Lakra, Mrs. Savita Patil, Mrs. Triparna Majumdar, Mr. Hitesh Dighe, Ms. Manisha Dudhmal, Mr. Yash, Mr. Vishwajit Dhanore from Maximum Containment Facility, ICMR-NIV, Pune. Authors would also like to acknowledge Dr. Priya Abraham, Director ICMR-NIV, Pune for her support.

