## Supplementary table 1, Supplementary table 2, Supplementary table 3, for "Isolation and characterization of SARS-CoV-2 VOC, 20H/501Y.V2, from UAE travelers"

**Supplementary table 1. Clinical details of the international traveler screened for SARS-CoV-2 at Delhi International Airport, India**

| **Sr.no.** | **Age range** | **Clinical presentation** | **States and number of cases** | **Date of arrival in India and sample Collection** | **International Travel History and number of cases** | **SARS-CoV-2 result with Ct Value**  **(E gene) range** | **SARS CoV-2 result with Ct Value (RdRp2) range** |
| --- | --- | --- | --- | --- | --- | --- | --- |
| 1 | 24-68 years | Asymptomatic | Uttar Pradesh (n=13), Bihar (n=9), Punjab (n=6), Himachal Pradesh (n=2), Jammu Kashmir (n=2), New Delhi (n=2), Gujarat (n=1), Haryana (n=1), Odisha (n=1), Rajasthan (n=1), West Bengal (n=1) | 24 February 2021-  16 March 2021 | UAE (n=39) | 16.8-29.7 | 16.5-30.8 |
| 2 | 24-48 years | Asymptomatic | Punjab (n=3), Uttar Pradesh (n=2), New Delhi (n=2), Chhattisgarh (n=1), Kerala (n=1), Uttarakhand (n=1) | 23 February 2021-  5 March 2021 | West and East Africa (n=10) | 17-29.5 | 15.9-28.7 |
| 3 | 26-49 years | Asymptomatic | West Bengal (n=3), Punjab (n=1), Bihar (n=1) | 10 March 2021-  16 March 2021 | Qatar (n=5) | 24.4-30.4 | 24-29 |
| 4 | 24-27 years | Asymptomatic | Uttar Pradesh (n=1), Bihar (n=1), Odisha (n=1) | 5 March 2021-  6 March 2021 | Ukraine (n=3) | 16-19-9 | 16.1-23.8 |
| 5 | 27 years | Asymptomatic | Odisha (n=1) | 10 March 2021 | Saudi Arabia (n=1) | 22.4 | 23.4 |

*Ct= Cycle threshold, RdRP=RNA dependent RNA polymerase, UAE=United Arab Emirates

Supplementary table 2. B.1.351 SARS-CoV-2 isolates nucleotide changes in different passages of VeroCCL81 with respect to the Wuhan HU-1 isolate.

| **Reference Genomic Position** | | | | | | | **Reference nucleotide** | **Allele nucleotide** | **Amino acid change** | **Overlapping annotations** |
| --- | --- | --- | --- | --- | --- | --- | --- | --- | --- | --- |
| **MCL-21-H-893** | | **MCL-21-H-1093** | | **MCL-21-H-1195** | **MCL-21-H-1439** | |  |  |  |  |
| **P2** | **P3** | **P2** | **P3** | **P2** | **P2** | **P3** |  |  |  |  |
| 174 | 174 | 174 | 174 | 174 | 174 | 174 | G | T |  | 5' UTR |
| 241 | 241 | 241 | 241 | 241 | 241 | 241 | C | T |  |  |
|  |  |  |  |  | 503 | 503 | C | T | Pro80Ser | ORF1ab |
| 1059 | 1059 | 1059 | 1059 | 1059 | 1059 | 1059 | C | T | Thr265Ile |  |
|  |  |  |  | 1714 |  |  | C | T |  |  |
|  |  | 2692 |  |  | 2692 | 2692 | A | T |  |  |
| 3037 | 3037 | 3037 | 3037 | 3037 | 3037 | 3037 | C | T |  |  |
| 4897 | 4897 |  |  |  |  |  | C | T |  |  |
|  |  | 5100 | 5100 |  | 5100 | 5100 | C | T | Ser1612Leu |  |
|  |  |  |  | 5213 |  |  | T | C |  |  |
| 5230 | 5230 | 5230 | 5230 | 5230 | 5230 | 5230 | G | T | Lys1655Asn |  |
|  |  |  |  | 5371 |  |  | A | G |  |  |
| 5512 | 5512 |  |  |  |  |  | C | T |  |  |
| 6471 | 6471 |  |  |  |  |  | C | T | Thr2069Ile |  |
| 6762 | 6762 |  |  |  |  |  | C | T | Thr2166Ile |  |
|  |  |  |  | 10116 |  |  | C | T | Thr3284Ile |  |
| 10323 | 10323 | 10323 | 10323 | 10323 | 10323 | 10323 | A | G | Lys3353Arg |  |
|  |  |  |  | 10341 |  |  | C | T | Pro3359Leu |  |
|  |  |  |  |  | 10761 | 10761 | A | G | Lys3499Arg |  |
| 10809 | 10809 |  |  |  |  |  | C | T | Pro3515Leu |  |
| 11288 | 11288 | 11288 | 11288 | 11288 | 11288 | 11288 | TCTGGTTTT | - | Ser3675_Phe3677del |  |
|  |  |  |  | 11477 |  |  | A | C | Ile3738Leu |  |
|  |  | 12357 | 12357 |  |  |  | C | T | Thr4031Ile |  |
| 13059 | 13059 |  |  |  |  |  | C | T | Thr4265Ile |  |
| 14408 | 14408 | 14408 | 14408 | 14408 | 14408 | 14408 | C | T | Pro4715Leu |  |
|  |  |  |  | 16086 |  |  | T | C |  |  |
|  |  | 17733 | 17733 |  |  |  | C | T |  |  |
| 17999 | 17999 |  |  | 17999 |  |  | C | T | Thr5912Ile |  |
| 18657 | 18657 |  |  |  |  |  | C | T |  |  |
|  |  |  |  | 19803 |  |  | C | T |  |  |
|  |  |  |  | 21110 |  |  | C | T |  |  |
| 21077 | 21077 |  |  |  |  |  | C | T | Thr6938Ile |  |
|  |  |  |  | 21635 |  |  | C | T |  |  |
| 21801 | 21801 | 21801 | 21801 | 21801 | 21801 | 21801 | A | C | Asp80Ala | S |
|  |  |  |  |  | 22201 | 22201 | G | T |  |  |
| 22206 | 22206 | 22206 | 22206 | 22206 | 22206 | 22206 | A | G | Asp215Gly |  |
| 22281 | 22281 | 22281 | 22281 | 22281 | 22281 | 22281 | CTTTACTTG | - | Leu242_Leu244del |  |
| 22813 | 22813 | 22813 | 22813 | 22813 | 22813 | 22813 | G | T | Lys417Asn |  |
| 23012 | 23012 | 23012 | 23012 | 23012 | 23012 | 23012 | G | A | Glu484Lys |  |
| 23063 | 23063 | 23063 | 23063 | 23063 | 23063 | 23063 | A | T | Asn501Tyr |  |
| 23403 | 23403 | 23403 | 23403 | 23403 | 23403 | 23403 | A | G | Asp614Gly |  |
| 23664 | 23664 | 23664 | 23664 | 23664 | 23664 | 23664 | C | T | Ala701Val |  |
|  |  |  |  | 25546 |  |  | C | T | Leu52Phe | ORF3a |
| 25563 | 25563 | 25563 | 25563 | 25563 | 25563 | 25563 | G | T | Gln57His |  |
| 25904 | 25904 | 25904 | 25904 | 25904 | 25904 | 25904 | C | T | Ser171Leu |  |
| 26456 | 26456 | 26456 | 26456 | 26456 | 26456 | 26456 | C | T | Pro71Leu | E |
|  |  |  |  | 27059 |  |  | C | T |  | M |
|  |  | 27870 | 27870 |  | 27870 | 27870 | G | T | Glu39* | CDS: ORF7b |
|  |  |  |  | 27987 |  |  | G | T | Val32Leu | CDS: ORF8 |
|  |  | 28079 | 28079 |  |  |  | G | T |  |  |
| 28250 |  |  |  |  |  |  |  |  |  |  |
|  | 28253 | 28253 | 28253 | 28253 | 28253 | 28253 | CA | TC | Ile121Leu |  |
| 28254 | 28254 |  |  |  |  |  | A | - | Ile121fs |  |
| 28887 | 28887 | 28887 | 28887 | 28887 | 28887 | 28887 | C | T | Thr205Ile | CDS: N |
|  |  | 29358 | 29358 |  | 29358 | 29358 | C | T | Thr362Ile |  |
|  |  | 29555 | 29555 |  |  |  | C | T |  |  |

Supplementary table 3. The percent nucleotide similarity of representative B.1.351 sequences from GISAID respect to B.1.351 isolates from India and Wuhan-Hu-1

|  | **NC_045512.2 isolate_Wuhan-Hu-1** | **MCL-21-H-1195** | **MCL-21-H-893** | **MCL-21-H-1439** | **MCL-21-H-1093** |
| --- | --- | --- | --- | --- | --- |
| hCoV-19/South_Africa/MCL-21-H-1195/2021/ EPI_ISL_2014135 | 99.89 |  |  |  |  |
| hCoV-19/South_Africa/MCL-21-H-893/2021/ EPI_ISL_2014131 | 99.90 | 99.93 |  |  |  |
| hCoV-19/South_Africa/MCL-21-H-1439 /2021/ EPI_ISL_2014133 | 99.91 | 99.93 | 99.94 |  |  |
| hCoV-19/South_Africa/MCL-21-H-1093/2021/ EPI_ISL_2014132 | 99.91 | 99.92 | 99.93 | 99.97 |  |
| hCoV-19/South_Africa/Tygerberg_650/2020\|EPI_ISL_1591449\| | 99.91 | 99.92 | 99.94 | 99.96 | 99.95 |
| hCoV-19/South_Africa/NHLS-UCT-GS-0821/2020\|EPI_ISL_960139\| | 99.92 | 99.93 | 99.94 | 99.96 | 99.95 |
| hCoV-19/South_Africa/NHLS-UCT-GS-0812/2020\|EPI_ISL_700551\| | 99.92 | 99.93 | 99.94 | 99.95 | 99.95 |
| hCoV-19/South_Africa/NHLS-UCT-GS-0808/2020\|EPI_ISL_700441\| | 99.92 | 99.95 | 99.96 | 99.96 | 99.96 |
| hCoV-19/South_Africa/NHLS-UCT-GS-0807/2020\|EPI_ISL_700492\| | 99.92 | 99.93 | 99.94 | 99.95 | 99.94 |
| hCoV-19/South_Africa/NHLS-UCT-GS-0731/2020\|EPI_ISL_960136\| | 99.93 | 99.96 | 99.97 | 99.97 | 99.97 |
| hCoV-19/South_Africa/NHLS-UCT-GS-0684/2020\|EPI_ISL_700535\| | 99.92 | 99.93 | 99.94 | 99.95 | 99.95 |
| hCoV-19/South_Africa/NHLS-UCT-GS-0683/2020\|EPI_ISL_700450\| | 99.92 | 99.94 | 99.95 | 99.97 | 99.96 |
| hCoV-19/South_Africa/NHLS-UCT-GS-0677/2020\|EPI_ISL_700469\| | 99.92 | 99.93 | 99.94 | 99.95 | 99.95 |
| hCoV-19/South_Africa/NHLS-UCT-GS-0676/2020\|EPI_ISL_700510\| | 99.92 | 99.93 | 99.94 | 99.95 | 99.95 |
| hCoV-19/South_Africa/NHLS-UCT-GS-0672/2020\|EPI_ISL_960129\| | 99.93 | 99.93 | 99.94 | 99.95 | 99.95 |
| hCoV-19/South_Africa/NHLS-UCT-GS-0671/2020\|EPI_ISL_700483\| | 99.92 | 99.93 | 99.94 | 99.96 | 99.95 |
| hCoV-19/South_Africa/NHLS-UCT-GS-0669/2020\|EPI_ISL_700482\| | 99.92 | 99.93 | 99.94 | 99.95 | 99.95 |
| hCoV-19/South_Africa/NHLS-UCT-GS-0666/2020\|EPI_ISL_700587\| | 99.92 | 99.93 | 99.94 | 99.95 | 99.95 |
| hCoV-19/South_Africa/NHLS-UCT-GS-0662/2020\|EPI_ISL_700460\| | 99.92 | 99.93 | 99.94 | 99.95 | 99.95 |
| hCoV-19/South_Africa/NHLS-UCT-GS-0660/2020\|EPI_ISL_700438\| | 99.91 | 99.92 | 99.93 | 99.94 | 99.94 |
| hCoV-19/South_Africa/NHLS-UCT-GS-0659/2020\|EPI_ISL_700581\| | 99.92 | 99.93 | 99.94 | 99.95 | 99.95 |
| hCoV-19/South_Africa/NHLS-UCT-GS-0645/2020\|EPI_ISL_700452\| | 99.92 | 99.95 | 99.96 | 99.96 | 99.96 |
| hCoV-19/South_Africa/NHLS-UCT-GS-0644/2020\|EPI_ISL_700451\| | 99.92 | 99.95 | 99.96 | 99.96 | 99.96 |
| hCoV-19/South_Africa/NHLS-UCT-GS-0634/2020\|EPI_ISL_960134\| | 99.92 | 99.93 | 99.94 | 99.95 | 99.95 |
| hCoV-19/South_Africa/NHLS-UCT-GS-0633/2020\|EPI_ISL_700478\| | 99.92 | 99.93 | 99.94 | 99.95 | 99.94 |
| hCoV-19/South_Africa/N00393/2020\|EPI_ISL_712094\| | 99.87 | 99.88 | 99.89 | 99.90 | 99.88 |
| hCoV-19/South_Africa/N00390/2020\|EPI_ISL_712081\| | 99.92 | 99.93 | 99.94 | 99.95 | 99.94 |
| hCoV-19/South_Africa/N00385/2020\|EPI_ISL_712073\| | 99.92 | 99.94 | 99.94 | 99.97 | 99.96 |
| hCoV-19/South_Africa/N00380/2020\|EPI_ISL_712071\| | 99.93 | 99.94 | 99.95 | 99.97 | 99.96 |
| hCoV-19/South_Africa/N00347/2020\|EPI_ISL_712079\| | 99.92 | 99.95 | 99.96 | 99.96 | 99.96 |
| hCoV-19/South_Africa/N00334/2020\|EPI_ISL_712086\| | 99.91 | 99.93 | 99.94 | 99.96 | 99.95 |
| hCoV-19/South_Africa/KRISP-K004644/2020\|EPI_ISL_660659\| | 99.93 | 99.95 | 99.96 | 99.97 | 99.97 |
| hCoV-19/South_Africa/KRISP-K004312/2020\|EPI_ISL_660190\| | 99.92 | 99.94 | 99.95 | 99.96 | 99.96 |
